# A non-vesicular Argonaute protein is transmitted from nematode to mouse and is important for parasite survival

**DOI:** 10.1101/2025.04.01.646544

**Authors:** Kyriaki Neophytou, Isaac Martínez-Ugalde, Thomas Fenton, Elaine Robertson, Lewis J. Strachan, Yvonne Harcus, Chanel M. Naar, David Wright, Daniel R. G. Price, Ruby White, Michael J. Evans, Jose R. Bermúdez-Barrientos, Hanchen Li, Rick M. Maizels, Raffi V. Aroian, Alasdair J. Nisbet, Cei Abreu-Goodger, Amy H. Buck

**Affiliations:** Institute of Immunology and Infection Research, School of Biological Sciences, The University of Edinburgh, Edinburgh, EH9 3FL, UK; Institute of Ecology and Evolution, School of Biological Sciences, The University of Edinburgh, Edinburgh, EH9 3FL, UK; Department of Molecular Genetics, Faculty of Medicine, University of Toronto, Toronto, ON M5S 1A8 Canada; School of Immunity & Infection, University of Glasgow, Glasgow, G12 8TA, UK; Cambridge Institute for Medical Research, University of Cambridge, Cambridge, CB2 0XY, UK; Leiden University Center for Infectious Diseases, Leiden University Medical Center, Leiden, 2333ZA, Netherlands; Department of Vaccines and Diagnostics, Moredun Research Institute, Edinburgh, EH26 0PZ, UK; Department of Disease Control, Moredun Research Institute, Edinburgh, EH26 0PZ, UK; Cancer Research UK Scotland Institute, Switchback Road, Bearsden, Glasgow, G61 1BD, UK; University of Massachusetts Chan Medical School, Program in Molecular Medicine, Worcester, MA, USA, 01605

**Keywords:** Argonaute, host-pathogen, extracellular RNA, RNA interference, nematode, parasite, RNA communication

## Abstract

Argonautes are ancient proteins with well-characterised functions in cell-autonomous gene regulation and genome defense but less clear roles in non-cell-autonomous processes. Extracellular Argonautes have been reported across plants, animals and protozoa yet their biochemical and functional properties remain elusive. Here we demonstrate that an extracellular Argonaute (exWAGO) released by the rodent-infective parasitic nematode *Heligmosomoides bakeri* is detectable inside mouse cells during the natural infection. We further show that exWAGO is released from *H.bakeri* in both vesicular and non-vesicular forms that have different resistances to proteolysis, different accessibilities to antibodies and associate with different subsets of secondary siRNAs. Using recombinant exWAGO protein we demonstrate that non-vesicular exWAGO is directly internalised by mouse cells *in vitro* and that immunisation of mice with exWAGO confers partial protection against subsequent *H. bakeri* infection and generates antibodies that block exWAGO uptake into cells. Finally, we show that properties of exWAGO are conserved across Clade V nematodes that infect humans and livestock. Together this work expands the context in which Argonautes function and illuminates an RNA-binding protein as a vaccine target for parasitic nematodes.

## Introduction

Argonautes are ancient proteins that work in partnership with small nucleic acid guides to regulate gene expression, control transposable elements and defend cells against viruses. In animals there are two highly conserved clades of Argonautes: the AGO family that acts with microRNAs (miRNAs) or short interfering RNAs (siRNAs), and the PIWI family that acts with PIWI RNAs (piRNAs) (Swarts *et al*, 2014). Nematodes have evolved an additional clade of Argonautes, the Worm-specific Argonautes (WAGOs), which bind secondary siRNAs that are generally 22 nucleotides in length, start with a Guanosine (G) nucleotide and commonly have a 5’ triphosphate moiety due to their generation by RNA-dependent RNA polymerases (Gu *et al*, 2009; Seroussi *et al*, 2023; Yigit *et al*, 2006). Research in *Caenorhabditis elegans* suggests that WAGOs function to amplify or transmit an RNA interference (RNAi) response, for example during environmental RNAi (amplifying responses against foreign RNA) or soma-to-germline transmission of RNAi (transgenerational inheritance) (Ketting & Cochella, 2020). However, many nematode species are parasitic and how they might use Argonautes during infections inside diverse hosts has not been well explored (Buck & Blaxter, 2013).

In order to survive in their hosts, parasitic nematodes release excretory-secretory (ES) products that condition the host environment and immune response towards tolerance, also promoting chronic infections (Maizels, 2020; Girgis *et al*, 2013; King & Li, 2018). *Heligmosomoides bakeri* is a gastrointestinal nematode parasite of mice that serves as a model for intestinal worm infections, which are estimated to affect a quarter of the world’s human population (WHO, 2023; Reynolds *et al*, 2012) and are highly prevalent in domestic and wild animal species, causing substantial global health burdens and economic losses (Charlier *et al*, 2020). We previously showed that extracellular vesicles (EVs) are a component of *H. bakeri* ES (HES) and demonstrated that the parasite EVs have immune suppressive properties in host cells (Buck *et al*, 2014; Coakley *et al*, 2017). We subsequently discovered that a specific WAGO protein (Hb-exWAGO) is present in these EVs in association with siRNAs (Chow *et al*, 2019). Yet whether EVs are the main carrier of siRNAs and exWAGO is not clear, nor is it known whether Hb-exWAGO serves a function during infection. Studies in plants have demonstrated that small RNA trafficking from fungal or oomycete pathogens into plant cells is important for pathogen survival and immune evasion (Qiao *et al*, 2023). Yet in these systems the pathogen small RNAs hijack the Argonaute of the receiving host cell to function (Qiao *et al*, 2023). There is no example to date of an Argonaute from one species operating in another, despite a mounting body of literature across plants, protozoa and animals that Argonautes can get out of cells in multiple forms (He *et al*, 2021; Geekiyanage *et al*, 2020; Arroyo *et al*, 2011; McKenzie *et al*, 2016; Melo *et al*, 2014; Karimi *et al*, 2022; Si *et al*, 2023; Chow *et al*, 2019; Garcia-Silva *et al*, 2013; Zhang *et al*, 2019, 2021; Jeppesen *et al*, 2019).

Here we take advantage of vaccination as a method to target the environment-exposed form of Hb-exWAGO in *H. bakeri*. We show that vaccination with recombinant Hb-exWAGO significantly reduces parasite burden and that antibodies raised during infection block the uptake of non-vesicular Hb-exWAGO into cells *in vitro.* We also provide evidence of transfer of the Hb-exWAGO from nematode to mouse cells during the natural infection using immunohistochemistry. Finally, we show that the properties of exWAGO are conserved in Clade V parasitic nematodes that infect humans and livestock, including high expression in adult stages, presence in ES products and preference for binding secondary siRNAs. This work illuminates the functional capacity of an extracellular Argonaute and reveals a new vaccine candidate for an important and neglected class of pathogens where there are no vaccines for humans (Perera & Ndao, 2021), limited (and no recombinant) vaccine options in animals (Claerebout & Geldhof, 2020) and mounting resistance to anthelminthic drugs (Orr *et al*, 2019; Wit *et al*, 2021).

## Results

### Non-vesicular and vesicular Hb-exWAGO both bind to secondary siRNAs but have different accessibilities to proteases and antibodies

In order to isolate vesicular and non-vesicular forms of Hb-exWAGO we used sequential centrifugation, filtration and ultracentrifugation of *H.bakeri* excretory-secretory products (HES). Using western blot analysis, we found that Hb-exWAGO in EVs was largely protected from proteolysis following proteinase K incubation in the absence of detergent whereas Hb-exWAGO in EV-depleted HES was susceptible (Figure 1A). We then developed an ELISA assay to test whether vesicular and non-vesicular Hb-exWAGO had different accessibilities to antibody capture. Using equal total protein inputs from EVs and EV-depleted HES, we detected an increase in the Hb-exWAGO signal when EVs were treated with detergent, whereas the Hb-exWAGO signal in EV-depleted HES remained unaffected (Figure 1B). These data collectively demonstrate that Hb-exWAGO is present both inside and outside EVs with each form having different protease sensitivities and antibody accessibilities. Using the ELISA assay in the presence of detergent with a recombinant Hb-exWAGO standard we further quantified that approximately 8.2±1.0x10^6^ (mean ± S.E.M) total copies of Hb-exWAGO are secreted per adult worm per day *ex vivo* (Figure 1C). We note that EVs accounted for <5% of the total amount of protein found in HES products when purified by ultracentrifugation (Figure S1A) and western blot analysis confirmed that substantially more non-vesicular Hb-exWAGO is released by the parasites compared to vesicular Hb-exWAGO (Figures 1D and S1B). We then carried out immunoprecipitation of Hb-exWAGO to determine whether both the vesicular and non-vesicular forms bind small RNAs. As shown in Figure 1E (upper panel), Hb-exWAGO isolated from both EVs and EV-depleted HES binds to small RNAs that are largely 22-23 nucleotides (nt) in length and start with a Guanosine, features that are consistent with secondary siRNAs (Chow *et al*, 2019). Consistent with our previous results using qRT-PCR of small RNAs bound to Hb-exWAGO in EVs, the majority of Hb-exWAGO-bound siRNAs in both EVs and EV-depleted HES derive from transposable elements (Chow *et al*, 2019) (Figure 1E lower panel). Strikingly however, multidimensional scaling (MDS) analysis shows that distinct siRNA populations are bound to Hb-exWAGO in EVs and EV-depleted HES, and both of these are also distinct from the total population of siRNAs bound to Hb-exWAGO within the adult worms (Figure 1F). Differential expression analysis of the genomic regions producing the Hb-exWAGO-bound siRNAs indicates that 45,379 genomic regions produce more siRNAs in the EV-depleted HES, while 16,194 regions produce more in EVs (Figure 1G). Collectively, these data demonstrate that Hb-exWAGO exists in both vesicular and non-vesicular forms, that these are not contaminants of one another, and that both forms are released in complex with distinct populations of siRNAs.

**Figure 1.**
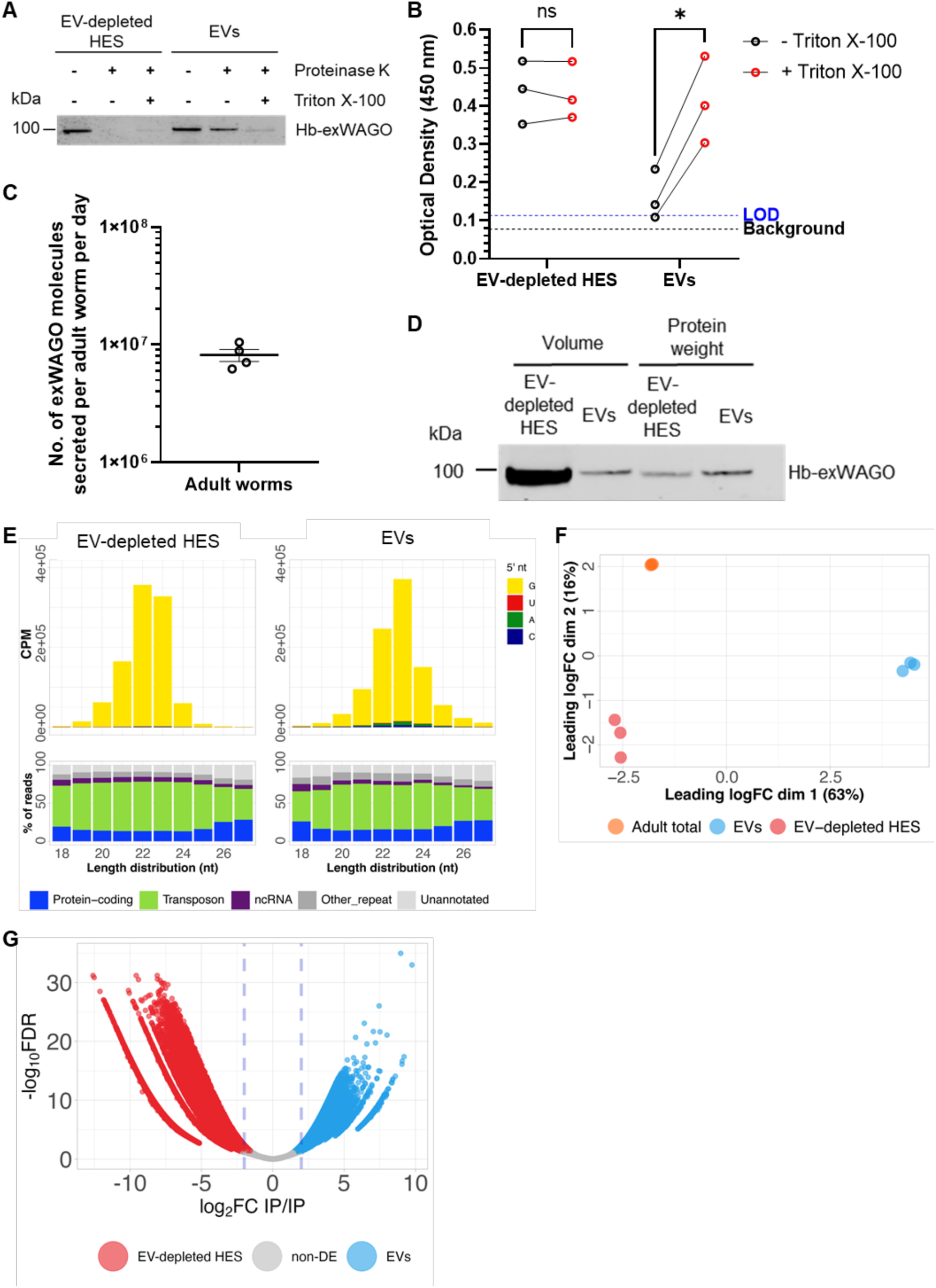
Hb-exWAGO is released in distinct vesicular and non-vesicular forms at high copy numbers. A) Western blot of Hb-exWAGO in EV-depleted HES or EVs following incubation in the absence or presence of Proteinase K (5 µg/ml) and/or Triton X-100 (0.05%) as indicated with 2.0 µg total protein. B) ELISA detection of Hb-exWAGO using equal protein amounts (0.2 µg) of EV-depleted HES or EVs in the presence or absence of Triton X-100 (0.05%) (n = 3 biological replicates). The limit of detection (LOD; blue dotted line) and background (no sample, black dotted line) are indicated. Significance was determined using a two-way ANOVA (ns = p > 0.05; * = p ≤ 0.05). C) Absolute copy number of Hb-exWAGO molecules present in HES, normalised per adult worm per day in culture based on sample of 30 worms (mixed sex) in culture from 24-96 hours post-harvest from mice. Data represent mean ± S.E.M (n = 4 biological replicates). D) Western blot of Hb-exWAGO in EV-depleted HES and EVs following separation by ultracentrifugation from the same volume of HES starting material (protein yield from EV-depleted HES = 38.5 µg, EVs = 2.0 µg) or using equal protein quantities (2.0 µg of each). E) Top: Length distribution and first nucleotide plots of reads mapping to the *H. bakeri* genome following Hb-exWAGO immunoprecipitation from EV-depleted HES or EVs. Data show the average length distributions (top panels) from n = 3 (CPM = counts per million). Bottom: The percentage of small RNA reads that map to different annotated regions of the *H. bakeri* genome. F) Multidimensional scaling analysis (MDS) of Hb-exWAGO small RNAs in adult worms, EVs and EV-depleted HES. Each dot represents an immunoprecipitation sample (n = 3 biological replicates), and the distance between them represents their similarity based on small RNAs aligned to distinct genomic regions. FC = fold-change. G) Volcano plot showing 45,379 genomic regions enriched in EV-depleted HES Hb-exWAGO immunoprecipitations (red) and 16,194 enriched in EVs (blue) using fold-change (FC) > 2 and false discovery rate (FDR) <0.05 as cutoffs. Non-enriched (Non-DE) regions are shown in grey. Each dot represents a distinct, non-overlapping genomic region.

### Hb-exWAGO is detected inside host cells *in vivo* and internalised by live cells *in vitro*

The above data suggest the adult worms release large quantities of Hb-exWAGO *ex vivo*. To test whether Hb-exWAGO is also released *in vivo* we used immunohistochemistry of paraffin-embedded *H. bakeri*-infected mouse gut sections at 7 days post infection. As shown in Figure 2A, Hb-exWAGO is detected inside the parasite, and also inside host immune cells that surround the worm. These data show that substantial quantities of Hb-exWAGO are detected *in vivo* and inside host cells, but do not distinguish which form of Hb-exWAGO, vesicular and/or non-vesicular, is transmitted to host cells.

**Figure 2.**
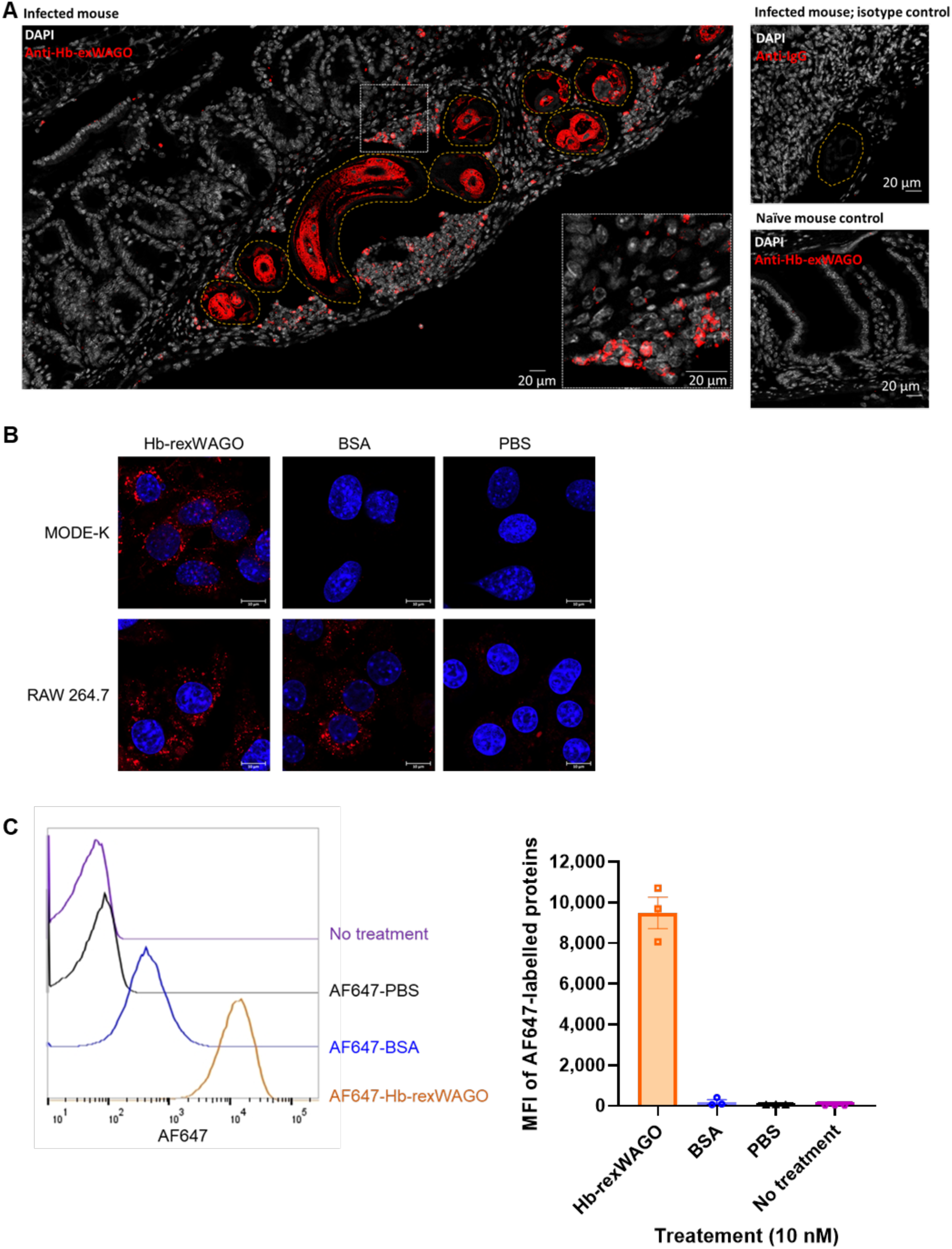
Detection of native Hb-exWAGO inside host cells *in vivo* and internalisation of recombinant Hb-exWAGO by live host cells *in vitro*. A) Confocal microscopy images of mouse gut tissue 7 days post infection with *H. bakeri* compared to uninfected gut tissue. Anti-Hb-exWAGO staining is shown in red. The outline of parasites is indicated with dotted lines. White box indicates zoomed in sections. Data are representative of 3 independent experiments. Scale bar = 20 µm. B) Confocal microscopy images of MODE-K and RAW 264.7 cells following incubation with 0.1 µM AF647-labelled recombinant Hb-exWAGO (Hb-rexWAGO), BSA or PBS for 4 hours. Scale bar = 10 µm. Data are representative of 3 independent experiments. C) The Median Fluorescence Intensity (MFI) of the AF647 signal detected by Flow cytometry analysis of MODE-K cells treated with 10 nM of AF647-labelled recombinant Hb-exWAGO (Hb-rexWAGO), BSA, PBS, or no treatment for 4 hours. Representative data are shown from 3 independent experiments (left) and data are from 3 independent experiments and show the mean ± S.E.M (right).

While we and others have shown uptake of parasite EVs by host cells, there is limited data on the capacity of non-vesicular nematode parasite proteins to enter live host cells, and no examples to date in this parasite model. We therefore first labelled EV-depleted HES with Cy-5 NHS Ester dye and tested whether EV-depleted HES proteins are internalised by mouse intestinal enterocyte epithelial MODE-K cells (Vidal *et al*, 1993). Signal was detected by 4 hours post incubation and remained strong at 24 hours with accumulation in the cytoplasm (Figure S2A). We then tested whether the recombinant Hb-exWAGO itself could be internalised, using BSA as a control for non-specific uptake. We compared uptake in the MODE-K cell line to a macrophage RAW 264.7 cell line expected to phagocytose extracellular material. As shown in Figure 2B, we detect internalisation of recombinant Hb-exWAGO by both MODE-K and RAW 264.7 cells, whereas BSA is only internalised by the RAW 264.7 cell line. We further developed a Flow Cytometry assay with MODE-K cells to measure recombinant Hb-exWAGO uptake down to picomolar concentrations (Figure S2B). As shown in Figure 2C, there is a clear shift in the Median Fluorescence Intensity in cells incubated with AF647-labelled Hb-exWAGO and near-background signal from the AF647-labelled BSA and PBS controls (Figures 2C and S2C). We note that cells were trypsinised to ensure any signal detected comes from internalised labelled proteins and not those that could be associated to the surface. Taken together, our results indicate that large quantities of Hb-exWAGO are released by adult parasites and are detected inside host cells *in vivo*, and the non-vesicular form of Hb-exWAGO is able to enter multiple cell types *in vitro* including epithelial cells.

### Vaccination with recombinant Hb-exWAGO confers partial protection against subsequent infection and generates antibodies that block Hb-exWAGO uptake into cells

To understand whether the non-vesicular form of Hb-exWAGO might be important for parasite survival in the host, we took advantage of the capacity to target this protein through vaccination with recombinant Hb-exWAGO. We used, as a benchmark, vaccination with total HES, which provides a cocktail of >360 proteins naturally released by the parasite (Buck *et al*, 2014; Hewitson *et al*, 2015). HES vaccination in mice was previously shown to result in sterile immunity with the observed effects attributed to both IgG1 antibody-mediated and Type 2 cellular responses (Hewitson *et al*, 2015).

Mice were immunised with recombinant Hb-exWAGO, PBS (negative control) or HES (positive control) in Imject Alum adjuvant followed by challenge with *H. bakeri* L3 stage larvae (Figure 3A). As expected from previous studies (Hewitson *et al*, 2015; Coakley *et al*, 2017), vaccination with HES confers sterile immunity as seen by a dramatic drop in the number of eggs per gram of faeces at 14 days post-challenge and clearance of worms by day 28 post-challenge compared to mice vaccinated with PBS (Figures 3B and C, and S3A). Strikingly, vaccination with recombinant Hb-exWAGO resulted in an average decrease of 67% in adult worm burdens compared to the PBS group at day 28 post-challenge (Figures 3B and S3A). Consistent with the reduction in worm burden, there were also decreased egg burdens, starting at day 14 post-challenge in the Hb-exWAGO-vaccinated group (Figure 3C). Serum antibody responses show that vaccination with recombinant Hb-exWAGO elicited IgG and, in particular, IgG1 antibodies specific to Hb-exWAGO (Figures 3D and S3B), consistent with known effects of alum adjuvant (Boyaka, 2017). Immunisation with HES also induced generation of Hb-exWAGO-specific IgG1 antibodies, as expected since Hb-exWAGO is present in HES, but these are much lower than in the Hb-exWAGO vaccine (Figure 3D). Interestingly, we detected Hb-exWAGO-specific IgG1 antibodies post-challenge in mice immunised with PBS indicating that (at least some of) the Hb-exWAGO protein becomes accessible to the immune system via the natural secretion of Hb-exWAGO by the parasites and that lab mice naturally generate a low level of Hb-exWAGO-specific antibodies during infection, but this is significantly boosted by the vaccine (Figure 3D).

**Figure 3:**
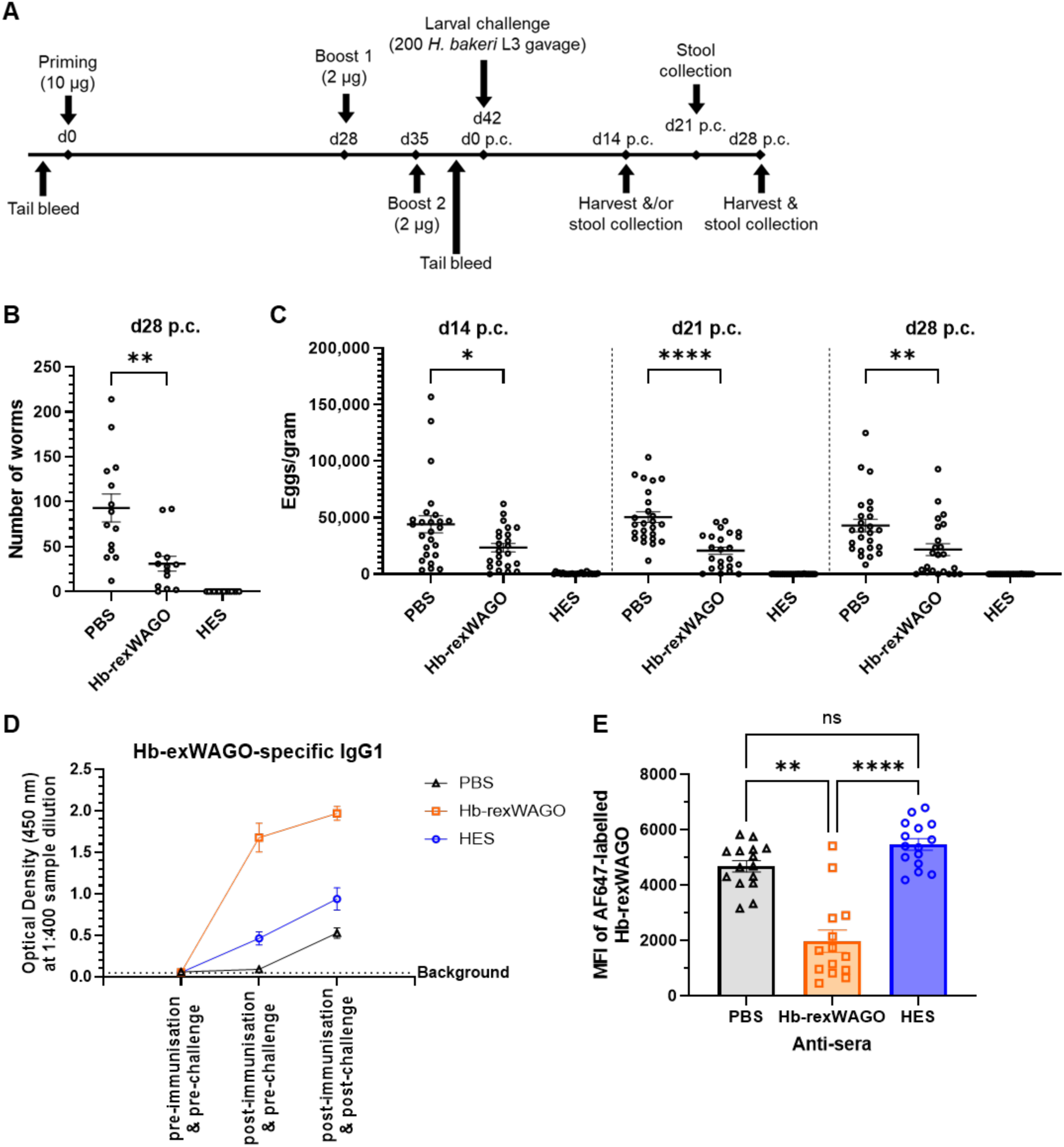
Vaccination with Hb-exWAGO generates partial protection against infection and generates antibodies that block Hb-exWAGO internalisation by mouse cells. A) Schematic of the immunisation timeline. Mice were vaccinated with recombinant Hb-exWAGO (Hb-rexWAGO), HES, or PBS and then challenged with 200 L3 stage *H. bakeri* larvae. B) The number of adult worms 28 days post-challenge (p.c.) recovered in the small intestine following vaccination of mice and challenge with 200 L3 stage larvae (n = 14 mice per vaccination group; details in SI Materials and Methods). C) The number of eggs per gram of faeces at 14, 21 and 28 days post-challenge (p.c.). (n = 23-25 mice per vaccination group). D) ELISA detection of exWAGO-specific IgG1 antibodies in sera obtained from vaccinated mice pre-immunisation and pre-challenge (pooled serum from one experiment), post-immunisation and pre-challenge, and post-immunisation and 28 days post-challenge (n = 5-14 mice per vaccination group). Data show the mean optical density at 1:400 sample dilution ± S.E.M. E) Median Fluorescence Intensity (MFI) of AF647-labelled Hb-exWAGO in MODE-K cells treated with 10 nM labelled protein for 4 hours and sera (1:250 dilution) from vaccinated mice 28 days post-challenge (n = 14-15 mice per vaccination group, from 3 independent experiments). All data are pooled from 3 independent experiments and represent mean ± S.E.M. B, C and E) Significance was determined using an unpaired Kruskal-Wallis test (ns = p > 0.05; * = p ≤ 0.05; ** = p ≤ 0.01, *** = p ≤ 0.001; ****= p ≤ 0.0001).

To test whether antibodies could inhibit Hb-exWAGO uptake by cells, we used the Flow Cytometry assay with labelled recombinant Hb-exWAGO. As shown in Figure 3E, treatment of MODE-K cells with sera from Hb-exWAGO vaccinated mice post-challenge inhibited uptake of AF647-labelled Hb-exWAGO compared to sera from mice vaccinated with HES and PBS. Reduction in the uptake of labelled Hb-exWAGO was also observed by confocal microscopy following treatment of MODE-K cells with anti-Hb-exWAGO antibodies compared to isotype controls (Figure S3C). These data suggest an antibody-exposed form of Hb-exWAGO is important for parasite survival *in vivo* and demonstrate that vaccination generates functional antibodies against Hb-exWAGO that block its uptake into host cells.

### ExWAGO is secreted across Clade V nematodes that infect livestock and humans and shows conserved features in its association with 22G RNAs

A notable feature of Hb-exWAGO is its high conservation across Clade V gastrointestinal nematodes that infect livestock and humans (spanning 65-81% amino acid identity, Dataset S1). To examine whether these exWAGO orthologues have similar properties to Hb-exWAGO, we first examined life stage expression of the exWAGO orthologues using publicly available transcriptome data from *H. bakeri*, rodent-infective *Nippostrongylus brasiliensis*, ruminant-infective *Teladorsagia circumcincta* and human-infective *Ancylostoma ceylanicum* nematodes. As shown in Figure 4A and Dataset S2, exWAGO expression is high in all the adult parasites, with lower levels in most larval stages across the four species. Using the ELISA assay, we also found that *H. bakeri* adults (d14 p.c.) release more Hb-exWAGO than larvae (d5 p.c.) and early adults (d7 p.c.) when using the same volume of ES collected from equal numbers of larvae or adults, as shown in Figure 4B. Western blot analysis of ES from *T. circumcincta* also showed that more exWAGO is found in ES from adults compared to larvae (Figure 4C). Furthermore, we detected exWAGO in the ES of the ruminant- and human-infective worm *Trichostrongylus colubriformis* and found it to be present in previously published proteomic analysis of the ES of these parasites (Figure 4C and Table S1). Collectively our data suggest that exWAGO is an abundant secreted protein expressed across the parasite developmental stages and is most highly expressed in adults.

**Figure 4:**
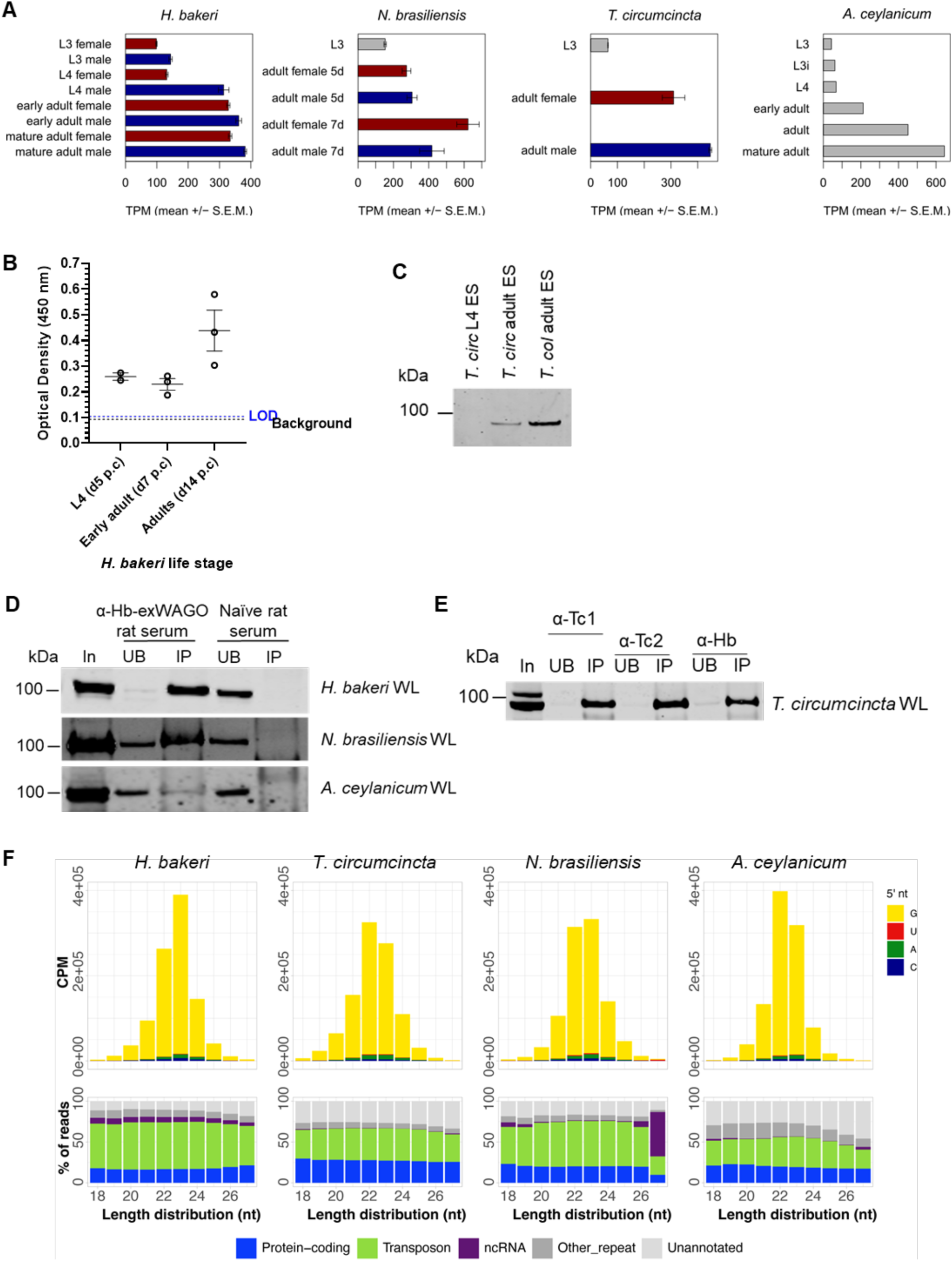
ExWAGO is highly expressed in adult worms across Clade V nematodes that infect livestock and humans and binds 22-23G small RNAs. A) Expression levels of exWAGO at different developmental stages based on RNAseq data in *H. bakeri* (PRJNA750155), *N. brasiliensis* (PRJNA994163; PRJEB20824), *T. circumcincta* (PRJEB7677) and *A. ceylanicum* (PRJNA231490). Grey = no sex defined; red = female; blue = male; TPM = transcripts per million. B) ELISA detection of Hb-exWAGO in HES collected from *H. bakeri* across different life stages in the presence of Triton X-100 detergent (0.05%). Data show the mean optical density ± S.E.M (n = 2-3 biological replicates). The Limit of Detection (LOD) (dotted blue line) and background (no sample, black dotted line) are indicated. C) Western blot analysis of exWAGO from the ES of *T. circumcincta* (*Tcirc*) L4 and adults, and *T. colubriformis* (*Tcol*) adults, probed using anti-*H. bakeri*-exWAGO antibody (6.0 µg lysate loaded per sample). D) Western blot of exWAGO immunoprecipitations (IPs) using rat anti-Hb-exWAGO serum or naïve rat serum from worm lysates (WL) of *H. bakeri* (200 µg), *N. brasiliensis* (25 µg) and *A. ceylanicum* (25 µg). For each IP the volumes loaded were as follows: *H. bakeri* input (In) = 1%, unbound (UB) = 1%, eluate (IP) = 5% by volume; *N. brasiliensis* and *A. ceylanicum* In = 32%, UB = 2%, IP = 37.5%. E) Western blot of exWAGO immunoprecipitations using rat anti-Tc-exWAGO (α-Tc1 or α-Tc2) or anti-Hb-exWAGO serum from *T. circumcincta* (150 µg) worm lysate (WL). (In = 6.7%, UB = 2%, IP = 10% by volume). F) Top: length distribution and first nucleotide (nt) plots of small RNA reads mapping to each parasite genome following exWAGO immunoprecipitation, polyphosphatase treatment and small RNA sequencing from *H. bakeri*, *N. brasiliensis*, *T. circumcincta* and *A. ceylanicum* adult worms. Data show the average length distributions (*H. bakeri*: n = 3; *N. brasiliensis*: n = 2, *T. circumcincta*: n = 2, and *A. ceylanicum*: n = 2) (CPM = counts per million). Bottom: percentage of reads that map to different annotated regions of each parasite genome.

Given the high sequence conservation of exWAGO orthologues in these parasite species, we tested whether antibodies generated against recombinant Hb-exWAGO could capture exWAGO in the other parasites. Using western blot analysis, we show that exWAGO was readily immunoprecipitated from *A. ceylanicum*, *N. brasiliensis* and *T. circumcincta* (Figures 4D and E). We then compared the profile of small RNAs bound to exWAGO in each species using small RNA sequencing and found that the small RNA profiles are extremely similar, showing the expected Guanosine as the first nucleotide, a length of 22-23 nucleotides and a propensity for mapping to transposable elements (Figure 4F). Comparison between polyphosphatase-treated and untreated libraries also showed an increase in the amount of small RNAs ligated to the adapters in response to 5’ RNA polyphosphatase treatment during the library preparation (Figure S4), consistent with the presence of a 5’ triphosphate. Taken together, these data strongly point towards functional conservation of exWAGO orthologues in Clade V parasitic nematodes.

## Discussion

This work provides the first evidence that an Argonaute protein (assumed mainly to operate inside of cells) is a tractable vaccine candidate and suggests the non-vesicular form of exWAGO is functionally relevant during nematode parasite infections. Vaccine development remains challenging for parasitic nematodes, given the complex life cycles of nematodes and the numerous mechanisms for immunomodulation that they have co-evolved with their hosts over millions of years (Perera & Ndao, 2021). There are currently only two licensed anti-nematode vaccines for animal use, both of which are limited in scalability and rely on native antigens or irradiated larvae (Claerebout & Geldhof, 2020); there are no licensed vaccines for humans despite growing resistance to anthelmintic drugs (Perera & Ndao, 2021). Work by ourselves and others has demonstrated that secreted RNAs are one type of immune evasion strategy used by nematodes (Buck *et al*, 2014; Liu *et al*, 2019; Tran *et al*, 2021; Ding *et al*, 2019) and we previously showed that Hb-exWAGO is released in complex with secreted siRNAs in EVs (Chow *et al*, 2019). Here we show a large proportion of Hb-exWAGO is released in a distinct non-vesicular form and that vaccination against this protein reduces subsequent burdens of infection (67% reduction in adult worm burdens) (Figure 3). As a single recombinant protein, this is substantial compared to existing vaccines which often use a cocktail of antigens (Britton *et al*, 2020), and its application could also be envisioned in multivalent strategies.

Our working hypothesis is that, by targeting Hb-exWAGO, we block a suite of small RNAs from entering and acting within host cells and we show here that antibodies generated by the vaccinations block recombinant Hb-exWAGO uptake by host cells *in vitro*. We cannot rule out that antibodies could also target a form of Hb-exWAGO that is within the parasite (for example if expressed and accessible to antibodies in the intestine) *in vivo*. However, using a quantitative assay, we demonstrate that substantial (approx. 10^7^) copies of Hb-exWAGO are released by each parasite per day (Figure 1C) and show with immunohistochemistry that Hb-exWAGO is clearly detected inside host cells during the infection (Figure 2). We do not yet know the precise cells that internalise exWAGO or whether Hb-exWAGO directly mediates gene silencing inside these cells. Hb-exWAGO does not contain the catalytic DEDD/H tetrad involved in RNA slicing and still little is known about the mechanism of gene silencing of the closest orthologs in *C. elegans*, SAGO-1, SAGO-2 and PPW-1 that share approximately 30% amino acid identity with Hb-exWAGO (Dataset S1). Based on its higher expression and release by adults, we speculate that Hb-exWAGO plays an important role in the chronic infection stage, but it also could act earlier.

Our work interfaces with an increasing number of reports suggesting extracellular Argonautes exist in multiple forms across plants and animals. In humans, these have been reported within EVs (Bukong *et al*, 2014; Mantel *et al*, 2016; McKenzie *et al*, 2016) as well as non-vesicular complexes including exomeres and supermeres (Jeppesen *et al*, 2019; Zhang *et al*, 2019, 2021), and other non-vesicular forms (Arroyo *et al*, 2011; Geekiyanage *et al*, 2020; Turchinovich *et al*, 2011). The dialogue around extracellular Argonautes has primarily focused on whether Argonautes are genuinely packaged within EVs or represent contaminants that co-purify (for example if associated with the surface of the EV) (Weaver & Patton, 2020). We address this point specifically here with assays showing that proteinase K sensitivity and antibody accessibility of the vesicular Hb-exWAGO occurs only in the presence of detergent (in contrast to the non-vesicular form) (Figure 1). We also show that the two forms of Hb-exWAGO bind to distinct subsets of secondary siRNAs. A key message from this study is that non-vesicular extracellular Argonautes (in addition to those within EVs) can also enter cells and this merits further understanding. The mechanism(s) by which non-vesicular Hb-exWAGO could enter cells remains largely unknown; there is only one study in this area that reports the human AGO2 can be internalised into recipient cells via the receptor neuropilin-1 (Prud’homme *et al*, 2016). Why the parasite releases two forms of Hb-exWAGO is not clear; we showed that Hb-exWAGO from EVs versus EV-depleted HES bind secondary siRNAs derived from different genomic locations. This suggests potentially different secretion origins within the parasite and also possibly different gene targets in the host. It is also possible that having two forms provides robustness to ensure some of the Hb-exWAGO will find its way into the host cell. Given the ubiquity of extracellular Argonautes including in humans, our work points to a need of better understanding the targeting properties and function of both vesicular and non-vesicular forms of Argonautes across living systems.

## Materials and Methods

### Ethics statement

All the mice used in this project were bred by the inhouse facilities at The University of Edinburgh. Experimental procedures were executed under a UK Home Office licence P635073CF as approved by qualified veterinarians and were carried out by qualified personnel in accordance with the UK Home Office guidelines.

### Recombinant exWAGO proteins

The recombinant *H. bakeri* exWAGO and *T. circumcincta* exWAGO used in this publication were produced by Sino Biological in insect cells and are 107.3 and 106.8 kDa respectively. The recombinant proteins were designed to have 3x FLAG tags (underlined) and a PreScission cleavage site (underlined and bold), followed by a 6x His tag (underlined and italics) at the N-terminus as shown below.

The recombinant Hb-exWAGO protein sequence is as follows: MDYKDDDDKDYKDDDDKDYKDDDDKAL**LEVLFQGP**ASG*HHHHHH*SGGGGSMDQLK TGMGQLSVGAVALPEKRSPGGIGNKVDFVTNLTELSLKPNVPYYKYDIRMYIVYKGNDALEHLKELTKQTKDDFPEQERKSAAVAVYKHLCKTYKDVFLPDGALLYDRAAVLFSAQRQLKLDGEEKQFMLPASVVSSAGPDATGIRVVIKKVKDQFQVTSNDLSKAVNVRDMERDKGILEVLNLAVSQKGYMETSQFVTYGSGVHYLFDHRALGFRDNELPELMDGKYMGIGLTKSVKVLEGDSGKGNSAFVVTDVTKGAFHVDEQNLMEKISQMSIFFDQRTGQSSFNAKNAMQPFNQKAILQQIKGLYVRTTYGKKKTFPIGNLAAAANALKFQTADGAQCTVEQYFKKHYNIQLKYPGMFTVSERHNPHTYYPVELLTVAPSQRVTLQQQTPDQVASMIKASATLPQTRLHQTKIMKDALDITPRNHNLATAGISVANGFTAVSGRVLPSPRIAYGGNQILRPVDNCKWNGDRSVFLEPAKLTNWAVCVTLTQQDARRLQIKEYISRVEMRCRNRGMQVDPVAEVFTLKHQTFDGLKEWYASQKQKNRRYLMFITSDGIKQHDSIKLLEVEYQIVSQEIKGSKVDAVVTKNQNQTLDNVVAKINMKLGGVNYNVMLGVKNDDKAFSWLNDKDRMFVGFEISNPPALSKVEIERGASYKMPSVLGWGANCAGNHQQYIGDYVYIQPRQSDMMGAKLSELIVDILKRFRAATTIAPRHIVLYFSGISEGQFSLVTDTYMRAVNTGIASLSPNYKPSVTAVAVSKDHNERIYKTNISGNRATEQNIPPGTVIDTKIVSPVINEFYLNSHSAFQGTAKTPKYSLLADNSKIPLDVIEGMTHGLCYLHEIVTSTVSVPVPLIVADRCAKRGHNVYIANSNQGEHSVNTIDEANAKLVNDGDLKKVRYNA.

The recombinant *T. circumcincta* exWAGO protein sequence is as follows: M<u>DYKDDDDKDYKDDDDKDYKDDDDK</u>AL**<u>LEVLFQGP</u>**ASG*<u>HHHHHH</u>*SGGGGSMADQL SGGMGKLSVAAVALPEKRAPGSLGTKLDFVTNLTGIKLKPNVPYYKYDVRMYIVYKGNDGREVLKELTKQTKDDFPEQERKMAAVAIYKHLVKSYKDIFPQDGQFFYDRAAVLFSAQREMKLGGPEKVITLPASLSPTAGSDAAGIRVVIKKVTDGYQVTSNDLMKAVNVRDCERDKGILEVLNLAVSQKGYMETSQFVTYGTGVHYLYDHRALGFRDNELPDLMDGKYMGIGLTKAVKVLEGDQGKSASAFVVTDVTKGAFHIDEQNLLEKISQMSIFFDPRTGQSTFSVKAAMQPHNMKSILQLIKGLYVRTTYGRKRTFPIGNLAAAPNALKLQTSDGVQCTIEQYFKKQYNVQLKYPGLFTVSERHNPHNYYPVELLTVAPSQRVTLQQQTPDQVASMIKASATLPSNRLHQTKVMKEALDITPRNAKLASAGINVEDGFTTVPGRVLPTPTILYGGSQTLKPVDNCKWNGDRSRFLEPAQLTNWAVCATLTQNDARRLQIKDYVARVESRCRAKGMQVEAAAEIFTLTKQNFDGLREFYAAQKKKNRKYLLFITSDGIKQHDLIKLLEVEYQIVSQEVKGSKVDSVMFKNQNQTLDNVIAKINMKLGGVNYNVVLGSKPNDPASKWLNDKDRLFVGFEISNPPALSKMEIERGATYKMPSVLGWGANCAANPQHYIGDYVYIKPRQSDMMGAKLSELIVEILKKFRGATSLAPRHIVLYFSGISEGQFSLVTDTYMKAINTGITSLSANYRPSVTALAVSKDHNERLYKSNISGSRANEQNIPPGSVVDTKIVSPVINEFYLNSHSAFQGTAKTPKYSLLADDSKIPLDVIEGMTHGLCYLHEIVTSTVSVPVPLIVADRCAKRGHNIFIANSNLGSAAVSSIEEANEKLVNHGELEKVRYNA.

### Parasite life cycle and collection of excretory/secretory products *H. bakeri*

For maintaining the life cycle of *H. bakeri*, CBA × C57BL/6 F1 male mice were infected with 400 *H. bakeri* L3 stage larvae by oral gavage and adult worms were harvested from the small intestine 14 days post-challenge. Adult worms were extensively washed and cultivated in serum-free media (RPMI 1640 supplemented with 1.2% glucose, 5 U/ml penicillin, 5 µg/ml streptomycin, 2mM L-glutamine and 1% gentamycin) as described in Johnston et al, (2015). For collection of the excretory/secretory products from *H. bakeri* (HES), the conditioned medium was collected on day 4 and 8 post-culture, however, the first 24 hours of culture medium was first removed to reduce any potential host contamination. The HES collected was spun (300 rcf, 5 min, room temperature) to remove eggs, filtered (0.22 μm Millex-GP filter), and stored at -20°C until required. HES (∼50-60 ml per batch of HES; batch is defined as the parasites collected and pooled from one group of infected mice) was concentrated to <12.5 ml using a 5 kDa Vivaspin (3,000 rcf, 4°C) (GE Healthcare). Concentrated HES was then subjected to ultracentrifugation (100,000 rcf, 70 min, 4°C) in polyallomer centrifuge tubes (Beckman Coulter) in a SW40 Ti swinging-bucket rotor (Beckman Coulter). The supernatant (SUP, also known as “EV-depleted HES”) was collected and concentrated to a final volume of 0.4-1.0 ml after buffer exchange in > 40 ml of PBS using a 5 kDa Vivaspin (3,000 rpm, 4°C) (GE Healthcare). The EV pellet was washed twice with 12.5 ml of cold PBS and spun as before. PBS washes were discarded, and the EV pellet was resuspended in 120-150 μl PBS. The protein concentration of EVs and EV-depleted HES was measured using the Qubit Protein Assay kit (Thermo Fisher Scientific) and the material was stored at -80°C until required. The number of particles in the purified EVs was measured using the ZetaView Nanoparticle Tracking Analysis (Particle Metrix) as per manufacturer’s instructions.

### N. brasiliensis

The *N. brasiliensis* life cycle was maintained by infection of Sprague-Dawley male rats with 3,000 infective L3 larvae. Adult worms were obtained 8 days later from small intestinal tissues in a Baermann apparatus as described in Lawrence et al, (1996). Worms were washed extensively in PBS, and stored at -80°C until required.

### A. ceylanicum

*A. ceylanicum* hookworm life cycle was maintained in hamsters (Hu *et al*, 2012, 2018). All housing and care of laboratory animals used in the hookworm study conform to the National Institutes of Health (N.I.H., U.S.A.) Guide for the Care and Use of Laboratory Animals in Research (see 18-F22) and all requirements and all regulations issued by the United States Department of Agriculture (U.S.D.A.), including regulations implementing the Animal Welfare Act (P.L. 89–544) as amended (see 18-F23). Adult worms were collected from the gut of hamsters at day 20 post-infection. Following extensive washes with Hanks’ Balanced Salt Solution, worms were frozen at -80°C until used.

### T. circumcincta *and* T. colubriformis

For maintaining the life cycle of *T. circumcincta and T. colubriformis,* helminth-free lambs (<6 months old) were orally infected with L3 stage parasites. For generation of L4 and adult parasites animals were orally challenged with 150,000 or 20,000 L3 stage larvae, respectively. Parasites at L4 stage (7 days post-infection) and adult stage (28 days post-infection) were harvested from either the sheep gastric stomach (for *T. circumcincta*) or the first 3 meters of the small intestine (for *T. colubriformis*). Worms were washed three times in PBS and then cultured in RPMI 1640 medium (Thermo Fisher Scientific) containing 20 mM HEPES pH 7.5, 1% (*w*/*v*) d-glucose, 2 mM L-glutamine, 1000 U/ml penicillin, 1000 μg/ml streptomycin, 200 μg/ml gentamycin and 25 μg/ml amphotericin B, at 37°C in 5% CO_2_. Culture conditioned media were harvested every 24 hours and replaced with fresh media, with worms maintained in culture for 120 hours post collection. At each supernatant collection, parasite viability was confirmed on the basis of structural integrity and motility.

Following collection, the culture supernatants were clarified by centrifugation at 300 rcf for 10 min at 4°C, filtered through a 0.2-μm Minisart syringe filter (Sartorius) and stored at - 70°C. For processing of excretory-secretory (ES) products, culture supernatants from all collections were pooled then concentrated and buffer exchanged into PBS using 10-kDa MWCO Amicon Ultra-15 Centrifugal Filter Units (Millipore Sigma), following the manufacturer’s guidelines. Protein concentration was determined using the Pierce™ BCA protein assay (Thermo Fisher Scientific) with bovine serum albumin standards, and aliquots of ES products were stored at -70°C prior to use.

### Cell culture

The immortalised intestinal epithelial MODE-K cell line (derived from the small intestine of C3H/He female mice) was kindly provided by Dominique Kaiserlian (French Institute of Health and Medical Research) and were cultured at 37°C with 5% CO_2_ in DMEM medium supplemented with fetal bovine serum (10%), L-glutamine (1%), penicillin/streptomycin (1%), sodium pyruvate (1%) and non-essential amino acids (1%) (Vidal *et al*, 1993). The murine macrophage RAW 264.7 cell line (ATCC) was cultured at 37°C with 5% CO_2_ in DMEM medium supplemented with fetal bovine serum (10%), L-glutamine (1%), and penicillin/streptomycin (1%).

### Proteinase K treatment

*H. bakeri* EVs and EV-depleted HES (2.0 μg of protein) were treated with or without Triton X-100 treatment (0.05%, 30 min, on ice), followed by Proteinase K digestion (5 μg/ml, 30 min, 37^°^C) (Epicentre) as appropriate. The reaction was quenched by addition of phenylmethanesulfonyl fluoride (5 mM, 1h, on ice) (Sigma). Samples were then prepared for western blot analysis.

### Western Blot analysis

Protein samples were reduced and separated on 4-12% Bis-Tris NuPAGE SDS gels (Thermo Fisher Scientific). The separated proteins were transferred on an Immobilon-FL PVDF membrane (Millipore) by wet transfer (100V, 105 min). The membrane was blocked and then incubated with the primary antibody (4°C, overnight, on roller). For detection of Hb-exWAGO protein a polyclonal rabbit antibody generated and purified against the peptide TKQTKDDFPEQERK (Eurogentec) was used at 1:4,000 in 5% BSA/0.1% TBST (Chow *et al*, 2019), unless otherwise specified. For detection of exWAGO orthologues in ES from *T. circumcincta* and *T. colubriformis,* a mouse monoclonal antibody (clone C5F7) generated in house as described in Hewitson et al, (2011) against recombinant Hb-exWAGO was used at 1:2,000 in 3% milk/0.1% TBST. Following incubation with primary antibodies, the membrane was incubated (1h, room temperature) with goat anti-rabbit IgG DyLight 800 (Thermo Fisher Scientific, SA5-35571) or goat anti-mouse IgG AF680 secondary antibodies (Thermo Fisher Scientific, A21058). The signal was visualised using the Odyssey CLx imaging system (LICOR) and band intensities were quantified using ImageJ (Fiji) as described in Stael et al, (2022).

### ELISA for detection and quantification of native Hb-exWAGO protein

Secreted Hb-exWAGO from EVs, EV-depleted HES and total HES was measured using ELISA. Immuno plates (Thermo Fisher Scientific) were coated with polyclonal rat anti-Hb-exWAGO antiserum capture antibody (raised against recombinant Hb-exWAGO in-house) in 0.06 M carbonate buffer (4°C, overnight) and blocked (4% BSA/TBS, 37°C, 2h). Samples were lysed as required with 0.05% Triton X-100 (30 min, on ice), and plated (4°C, overnight) in dilution buffer (1% BSA/TBST). For detection, the polyclonal rabbit UE3R2 anti-exWAGO primary antibody (produced by Sino Biological against recombinant Hb-exWAGO) was used (5 μg/ml in dilution buffer, 2h, room temperature) followed by incubation with goat anti-rabbit IgG-HRP (Cell Signalling, 7024S) secondary antibody (1:2,000 dilution, 37°C, 1h, in the dark). The reaction was developed using TMB substrate buffer (SeraCare), stopped (10% phosphoric acid), and the optical density was read at 450 nm on the Varioskan Flash (Thermo Fisher Scientific). Samples were analysed in technical duplicates and wells with buffer only (no sample) were used to determine the background signal.

For quantification of Hb-exWAGO, a standard curve with recombinant Hb-exWAGO with 0.05% Triton X-100 in dilution buffer was prepared in a four-fold serial dilution (from 0.01907 pg to 20 ng), and analysed using a 4PL regression (GraphPad Prism). The amount of Hb-exWAGO was interpolated using the optical density values obtained in the ELISA assay. The limit of detection was calculated from 5 independent experiments as the mean of blanks + (standard deviation of blanks * 3.3) and the limit of quantification was calculated as the mean of blanks + (standard deviation of blanks * 10).

For detection of Hb-exWAGO secreted at different *H. bakeri* life stages, female C57BL/6 mice were infected with 200 L3 stage larvae by oral gavage. At day 5, 7 and 14 post-challenge mice were culled and larvae or adult worms were collected from the gut wall or lumen of the small intestine respectively. Larvae were washed 5x with Hanks’ Balanced Salt Solution supplemented with penicillin-streptomycin (1%). Adult worms were treated with gentamycin (1 mg/ml) in Hanks’ Balanced Salt Solution supplemented with penicillin streptomycin (1%). Worms were then visually inspected for motility and any obvious injuries and were cultivated in groups of 20 in a 48-well plate with 500 µl serum-free culture media. The first 24 hours of culture medium was removed and replenished. HES was collected and replenished 2 and 4 days after culture. For the ELISA assay, day 2 and 4 HES were pooled by equal volume and tested directly (i.e., no PBS exchange), while culture medium only samples were used as a negative control.

For quantification of Hb-exWAGO secreted by adult worms harvested at day 14 post infection: Worms were cultivated in groups of 30 (15 male and 15 female as determined by visual inspection) in a 24-well plate with 400 µl serum-free culture media. The first 24 hours of culture medium was removed, replenished, and HES was collected 4 days post culture. The HES collected was tested directly (i.e., no PBS exchange) by ELISA, and culture medium only samples were used as negative control.

### Uptake of non-vesicular proteins

To examine if non-vesicular proteins are internalised by epithelial cells *in vitro*, proteins in EV-depleted HES or PBS were labelled using a Cy5 NHS Ester dye (AAT Bioquest, 151) according to the manufacturer’s instructions at a 15:1 (dye to protein) molar ratio. Unconjugated dye was removed using the Zeba 7K MWCO spin columns (Thermo Fisher Scientific, 89882) as per manufacturer’s instructions. The labelled EV-depleted HES and PBS were stored (-80°C, dark) until required. MODE-K cells (1.5x10^5^) were seeded in a 24-well plate with 400 μl MODE-K media overnight. The cells were treated with Cy5-labelled EV-depleted HES at a final concentration of 10 μg/ml or 50 μg/ml or with an equivalent volume of Cy5-labelled PBS as a negative control for 4 or 24 hours. The cells were washed twice with PBS, supplemented with fresh media, and imaged on the EVOS M7000 Imaging system (Invitrogen) with the EVOS LED Light Cube Cy5 2.0 using the EVOS 60x objective, fluorite, LWD. Data were analysed on the EVOS Analysis software v1.5.1479.304.

### Immunohistochemistry

To detect Hb-exWAGO *in vivo*, C57BL/6 female mice were infected with 200 L3 stage larvae by oral gavage. At day 7 of infection, mice were culled and the upper half of the small intestine was removed for preparation into a gut roll. Fat was trimmed away from the gut wall and the intestines were rolled inside-out onto wooden skewers. The intestines were fixed for 30 minutes in 4% PFA (diluted from 16% methanol-free PFA, VWR). The intestines were cut lengthwise and rolled onto a new skewer as ‘gut rolls’, then incubated for a further 5 hours in fresh 4% PFA. The gut rolls were washed twice in PBS and then stored overnight in 70% ethanol before processing sequentially in 80% ethanol (30 min), 95% ethanol (30 min, twice), 100% ethanol (30 min, twice), xylene (30 min, twice), and paraffin wax (40 minutes at 60°C), before embedding in paraffin wax. 10 µm sections were cut from the resulting blocks, which were mounted on Superfrost Plus slides (VWR), dried overnight, baked for one hour at 60°C and stored at 4°C. Wax was removed for staining by washing sequentially in xylene (3 min, twice), xylene/ethanol mixed 1:1 (3 min), 100% ethanol (3 min, twice), 95% ethanol (3 min), 70% ethanol (3 min), 50% ethanol (3 min), before a final rinse with water. Antigen retrieval involved a 30-minute treatment in citrate buffer (Abcam) in a 98°C water bath, before a further 20-minute cool-down outside the water bath. Slides were washed in TBS and then blocked using TBS buffer with 1% BSA and 0.1% Tween-20. For Hb-exWAGO detection, 2.5 µg/mL of rabbit anti-Hb-exWAGO antibody UE3R2 or the polyclonal rabbit IgG isotype-matched control antibody (Thermo Fisher Scientific, 02-6102) were incubated on the slides overnight in TBS buffer with 0.1% Tween-20 at 4°C. After washing three times in TBS buffer with 0.1% Tween-20, Alexa Fluor 647 anti-rabbit secondary antibody (diluted 1:2,000 in TBS with 0.1% Tween-20) (Thermo Fisher Scientific, A21245) was incubated on the slides for two hours at room temperature. DAPI diluted 1:1,000 was added to the slides and washed off twice in TBS before mounting in ProLong Gold (Thermo Fisher Scientific). The slides were imaged on a Zeiss LSM 980 microscope with Airyscan 2 detector, using the Zeiss 20x 0.8 NA Air objective, with 405 and 639 nm lasers. Data were acquired using the Zen Blue 3.5 (Zeiss) and processed using Zen Blue 3.3 software (Zeiss).

### Uptake and blockade of recombinant Hb-exWAGO

To test whether non-vesicular Hb-exWAGO is internalised by epithelial and macrophage cells *in vitro*, equimolar amounts of recombinant Hb-exWAGO, BSA and PBS were labelled with AF647 NHS Ester dye (Thermo Fisher Scientific, A37573) according to the manufacturer’s instructions at a 15:1 (dye to protein) molar ratio. Unconjugated dye was removed using the Zeba 7K MWCO spin columns (Thermo Fisher Scientific, 89882) as per manufacturer’s instructions. The labelled proteins and PBS were stored (-80°C, dark) until required.

To assess the uptake of AF647-labelled Hb-exWAGO by mouse cells using confocal microscopy, 1.0x10^4^ MODE-K or RAW 264.7 cells were seeded in an 18-well ibidi-treat µ-slide (Ibidi, 81816) in 100 µl culture medium overnight. The cells were incubated (37°C with 5% CO_2_) with 0.1 µM Hb-exWAGO, BSA, or PBS labelled with AF647 NHS Ester for 4 hours. For blocking the uptake of Hb-exWAGO in MODE-K cells, anti-Hb-exWAGO rat serum (produced in-house against recombinant Hb-exWAGO as described in Chow et al. (Chow *et al*, 2019)) or naïve rat serum were added at a 1:167 final dilution mixed with the labelled protein. Following incubation, the cells were washed twice with PBS, fixed with 4% PFA (10 min), washed again, stained with DAPI, washed again, and mounted using the ibidi mount medium (Ibidi, 50001). Slides were stored (4°C, dark) and imaged within three days of preparation. The samples were illuminated with 405 and 639 nm lasers and imaged on the Zeiss laser scanning confocal microscope LSM 980 with Airyscan 2 detector using the Zeiss 63x 1.4 NA Oil objective. Data were acquired using the Zen Blue 3.5 (Zeiss) and processed using the Zen Blue 3.3 software (Zeiss).

To assess the uptake and blocking of AF647-labelled recombinant Hb-exWAGO by mouse cells using flow cytometry, 0.8x10^5^ MODE-K cells were seeded in a 24-well plate in 500 µl culture medium overnight. The cells were incubated (37°C with 5% CO_2_) with 10 nM of AF674-labelled Hb-exWAGO, BSA, or PBS for 4 hours. For blocking the uptake of recombinant Hb-exWAGO in MODE-K cells, sera obtained from Hb-exWAGO-, HES-, or PBS-vaccinated mice were added to the cells at a 1:250 final dilution directly prior to addition of AF647-Hb-exWAGO. Following incubation, the cells were washed twice with PBS, trypsinised (5 min, at 37°C with 5% CO_2_), resuspended in 5% FBS/PBS and centrifuged (400 rcf, 5 min, room temperature). The cells were resuspended in 0.5% FBS/PBS and analysed on the MACSQuant Analyser 10 (Miltenyi Biotec) using the R1 channel. Data were analysed using FlowJo software v10.

### Immunisation of mice

For vaccine experiments, female C57BL/6 mice between 7-12 weeks old were immunised intraperitoneally with recombinant Hb-exWAGO or PBS or HES in Imject alum adjuvant (Thermo Fisher Scientific) at 1:1 volume ratio (total intraperitoneal injection volume was 200 μl). Mice were primed (10 μg) and boosted (2 μg) twice at 28 and 35 days post priming as described in Hewitson et al, (2015). 7 days following the second boost, the mice were challenged with 200 *H. bakeri* L3 stage larvae by oral gavage. Mice were culled 14 and 28 days post-challenge by CO_2_ overdose or by intraperitoneal administration with Dolethal (30 mg, Vetoquinol). We note that a prolonged period of 3 months between priming and the first boost occurred for one of the three independent experiments due to COVID-19 interruptions.

The small and large intestine were carefully removed, stretched and the number of adult worms counted. The number of eggs was measured using the McMaster Egg Counting Chamber (CellPath) following floatation with saturated sodium chloride solution as described in Camberis et al, (2003). The blood serum was acquired via tail bleed or for euthanised mice it was collected using cardiac puncture. The blood was incubated (1-2 hours, room temperature) and centrifuged (5,000 rcf, 5 min, room temperature). The supernatant containing the serum was centrifuged again as before and stored at -80°C until required.

### ELISA for measuring Hb-exWAGO-specific antibody responses

Hb-exWAGO-specific serum antibody responses were measured using an Enzyme-Linked Immunosorbent Assay (ELISA). Plates were coated with 0.05 μg of recombinant Hb-exWAGO (Sino Biological) in 0.06 M carbonate buffer (4°C, overnight), blocked (4% BSA/TBS, 37°C, 2h), and blood serum was plated in a two-fold serial dilution (4°C, overnight) in dilution buffer (1% BSA/TBST). The secondary antibodies goat anti-mouse IgG1-HRP (SouthernBiotech, 1070-05) or goat anti-mouse IgA-HRP (SouthernBiotech, 1040-05) were used (1:2,000 dilution, 37°C, 1h, in the dark). The reaction was developed using TMB substrate buffer (SeraCare), stopped (10% phosphoric acid), and the optical density was read at 450 nm on the Varioskan Flash (Thermo Fisher Scientific). Samples were analysed in technical duplicates and wells without sample were used to determine the background signal.

### exWAGO immunoprecipitation

exWAGO was immunopurified using polyclonal rat anti-exWAGO serum (generated in-house against the *H. bakeri* or *T. circumcincta* recombinant exWAGO protein) or naive rat serum antibodies conjugated to Protein G magnetic beads (Thermo Fisher Scientific, 10003D). In all immunoprecipitations, *H. bakeri* anti-exWAGO rat serum was used except for pull-downs from *T. circumcincta* worms where *T. circumcincta* anti-exWAGO rat serum was used as this was available. Beads were washed five times in cold Binding Wash Buffer (PBS, 0.02% Tween-20) and the antibodies were then conjugated on the washed beads by incubation (2h, 4°C, rotating wheel). Unconjugated antibody was then removed, and the beads were equilibrated with the appropriate lysis buffer three times.

To immunoprecipitate exWAGO and its orthologues from adult worms, nematodes were lysed by bead beating (5 mm steel bead, Qiagen) using the Tissue Lyser II (Qiagen) in pre-cooled cartridges for 2 min at 30 Hz twice in worm lysis buffer (150 mM NaCl, 10 mM Tris.HCl, 0.5 mM EDTA, 0.5% NP40) containing protease and phosphatase inhibitors (1 tablet per 5 ml, Roche), and 200 U/ml RNase inhibitors (Promega). Unlysed material was removed by centrifugation (16,100 rcf, 10 min, 4°C). The supernatant containing lysed worms was quantified using the Qubit Protein Assay kit (Thermo Fisher Scientific). Per immunoprecipitation, 150 μg of adult worm lysate protein was used.

To identify the small RNAs associated with the vesicular and non-vesicular Hb-exWAGO, we immunoprecipitated Hb-exWAGO from EV and EV-depleted HES derived from corresponding batches, using 10X more EVs by protein weight to account for lower amounts of exWAGO in EVs compared to EV-depleted HES: EVs (17 μg of protein) or EV-depleted HES (170 μg of protein) were lysed (20 min, on ice) in TBS/0.05% Triton X-100 with protease inhibitors (1 tablet per 10 ml, Roche) and 200 U/ml RNase inhibitors (Promega).

The lysate was incubated with the antibody-conjugated beads (45 min, 4°C, rotating wheel). The beads were washed with cold Low Salt buffer (50 mM Tris.HCl pH 7.5, 300 mM NaCl, 5 mM MgCl_2_, 0.5% Triton X-100 and 2.5% glycerol), followed by two washes with cold High Salt buffer (50 mM Tris.HCl pH 7.5, 800 mM NaCl, 10 mM MgCl_2_, 0.5% Triton X-100 and 2.5% glycerol) for 5 min at 4°C, using a rotating wheel. The beads were washed once more with Low Salt buffer, followed by a cold PNK buffer wash (50 mM Tris.HCl pH 7.5, 50 mM NaCl, 10 mM MgCl_2_, 0.5% Triton X-100). For western blot analysis proteins were eluted in denaturing buffer whereas for small RNA sequencing analysis the RNA was eluted directly in Qiazol (700 μl, 5 min, room temperature) (Qiagen).

### Small RNA libraries

RNA in Qiazol (Qiagen) was spiked with 7 μl of 10 pM RT4 synthetic spike (CTTGCGCAGATAGTCGACACGA) and extracted using the miRNA Serum/Plasma kit (Qiagen) according to the manufacturer’s instructions. The purified RNA was treated with RNA 5’ Polyphosphatase (Lucigen) according to the manufacturer’s instructions. The reaction was terminated by ethanol precipitation (-70°C, overnight) and the precipitated RNA was resuspended in 2.5 μl of nuclease-free water and 2.5 μl of Buffer 1 (TriLink, L-3206). The small RNA libraries were prepared using the CleanTag Small RNA Library Preparation Kit (TriLink, L-3206) according to the manufacturer’s instructions using half reaction volumes. All libraries were generated using 1:12 dilution of 5’ and 3’ adapters with 20 amplification cycles. The profile of each small RNA library prior to sequencing was assessed using either Novex 6% TBE acrylamide gels (Thermo Fisher Scientific) or the High Sensitivity DNA Bioanalyser chip (Agilent). Pooled libraries were size selected (140-220 bp for libraries made from adult worms or 140-180 bp for EV and EV-depleted HES libraries) using gel purification to remove adapter dimers. The libraries were sequenced on the Illumina NextSeq 2000 platform by the Edinburgh Clinical Research Facility or on the Illumina NovaSeq 6000 by Edinburgh Genomics using single-end 100 base pair reads.

### Small RNA bioinformatic analysis

The raw sequencing data were checked using FastQC v.0.12.1 and MultiQC v.1.14 (Andrews, 2010; Ewels *et al*, 2016) with no issues identified. The 3’ adapter sequence (TGGAATTCTCGGGTGCCAAGG) was removed using the Reaper tool from the Kraken package (Davis *et al*, 2013), and 18-27 nt reads were retrieved using Pullseq (available at: https://github.com/bcthomas/pullseq). ShortStack v.3.8 (Johnson *et al*, 2016) was then used to align reads to their respective genome, allowing up to one mismatch and weighted assignment of multi mapping reads (--nohp --mincov 5 --pad 1 --mismatches 1 --mmap u--bowtie_m all --ranmax ’none’). Count matrices were prepared across all samples, as well as plots showing the length distribution and first nucleotide preference, using the 01.featureCounts.R, 02.get_fn_mtx.R and 03.fn_long_cpm.R R scripts (available at: https://github.com/imu93/exwago_vaccine). Count values were transformed to counts per million (CPM) using the cpm function from the edgeR R package (Robinson *et al*, 2010). The data are deposited to the NCBI Sequence Read Archive (SRA) under the BioProject ID PRJNA1200757.

### Differential expression analysis of Hb-exWAGO immunoprecipitation from EVs and EV-depleted HES

Count matrices were first filtered to remove regions with less than five CPMs using the filterByExpr function from the edgeR (Robinson *et al*, 2010). The Trimmed Mean of M-values (TMM) normalisation and glmFit test were then used to estimate normalisation factors and fit negative binomial generalised log-linear models, respectively. The glmTreat test was used to identify regions significantly enriched in each secreted Hb-exWAGO form (Fold-change threshold = 2).

### Identification of exWAGO orthologues and Protein sequence alignment

Given the release of new genome assemblies for Strongylida parasites since the curation of exWAGO annotations (Chow *et al*, 2019), gene and protein exWAGO annotations of each genome were updated using miniprot (Li, 2023). Where available, the curated exWAGO protein was used. In cases where no curated exWAGO for that species was available, the sequence of the most closely related species was used for gene and protein prediction (e.g. *T. colubriformis* and *T. circumcincta*). Amino acid identity was further calculated using BLASTp against Hb-exWAGO annotated in the nHp.2.0 assembly (Chow *et al*, 2019).

### RNA sequencing data analysis

RNA-seq data was downloaded from NCBI SRA for the following species: *H. bakeri* (PRJNA750155), *N. brasiliensis* (PRJNA994163 and PRJEB20824), *T. circumcincta* (PRJEB7677) and *A. ceylanicum* (PRJNA231490). FastQC v.0.12.1 and MultiQC v.1.14 were used for quality control (Ewels *et al*, 2016; Andrews, 2010). Although almost all libraries showed no quality issues, those corresponding to the PRJNA994163 *N. brasiliensis* showed high proportions (∼15%) of Clontech SMART Primer II sequence and Illumina universal adapter. Thus, for these libraries an extra trimming step was implemented with cutadapt v.4.9 and parameters -g ^AAGCAGTGGTATCAAC -G^AAGCAGTGGTATCAAC -m.

The selected RNA-seq reads were aligned to their respective genome using STAR v.2.7.10b with the gtf gene annotation files to define junctions (--sjdbGTFfile gtf_annotation_file --clip3pAdapterSeq AGATCGGAAGAGCACACGT AGATCGGAAGAGCGTCGTG --outFilterMismatchNmax 3 –outFilterScoreMinOverLread 0.3) (Dobin *et al*, 2013). After alignment, gene expression was quantified using the featureCounts function from the Rsubread R package (Liao *et al*, 2019) (allowMultiOverlap= TRUE, requireBothEndsMapped = TRUE, countChimericFragments = FALSE, useMetaFeatures=TRUE, fraction = TRUE). The resulting raw counts were normalized into Transcripts Per Million (TPM) using the convertCounts function from the DGEobj.utils R package (Thompson *et al*, 2022). Each biological replicate was treated individually. If more than one orthologue was identified, the gene locus with the highest mapped reads and highest percentage amino acid identity was used (Dataset S1 & Dataset S2).

### Statistical analysis

Data were checked for normality using the Shapiro-Wilk test. For detection of Hb-exWAGO via ELISA, optical density values from EVs and EV-depleted HES treated with and without detergent were normally distributed and were analysed using a two-way ANOVA with Šídák’s correction for selected comparisons. Worm counts and faecal egg counts data were not normally distributed and were analysed using an unpaired Kruskal-Wallis test using uncorrected Dunn’s test for selected comparisons. Labelled recombinant Hb-exWAGO uptake inhibition data were not normally distributed and were analysed using an unpaired Kruskal-Wallis test with Dunn’s multiple comparisons test. Data analyses were performed using GraphPad Prism software version 10.1.2. Significance is denoted using asterisks where p < 0.05.

## Acknowledgments

This work was funded by ERC grant 101002385 to AHB. KN was funded by the Wellcome Trust PhD Programme grant 108905/Z/15/Z. IMU was funded by the Consejo Nacional de Ciencia y Tecnología (CVU: 896776; Scholarship number: 765004) and the Darwin Trust of Edinburgh. This work was also supported by the Wellcome Trust grant 203149/A/16/Z to the Wellcome Centre for Cell Biology which hosts the Centre for Optical Instrumentation Laboratory (now known as the Light Microscopy Facility core) and we wish to thank this facility for their support in training. AJN and DRGP are funded through the Strategic Research Programme (2022-2027) of the Scottish Government’s Rural and Environment Science and Analytical Services Division (RESAS). The *A. ceylanicum* work was supported by the National Institutes of Health - Eunice Kennedy Shriver National Institute of Child Health & Human Development grant 1R01-HD099072 to RVA. We thank the Bioresearch and Veterinary Services staff and facilities in King’s Buildings at The University of Edinburgh, Nicola Logan, Xiaochen Du, Rowan Bancroft and Sam Hillman for their support with vaccine experiments. We also thank Amy Pedersen and Matthew Taylor for discussions on our initial vaccination results and Tovah Shaw for sharing reagents and knowledge on immunohistochemistry. We also appreciate Franklin W. N. Chow for discussion and initial design/testing of Hb-exWAGO antibodies.

## Author contributions

K.N., I.M.U., T.F., C.A.G. and A.H.B. designed research; K.N., I.M.U., T.F., E.R., L.S., Y.H., C.M.N., D.W., D.R.G.P., R.W., M.J.E., J.R.B.B., H.L. and C.A.G. performed research; K.N., D.W., D.R.G.P., H.L., R.M.M., R.V.A., A.J.N. contributed new reagents/analytic tools; K.N., I.M.U., T.F., E.R., L.S., C.M.N., D.W., J.R.B.B. and C.A.G. analysed data, K.N. and A.H.B. wrote the paper.

## Resource availability

**Lead contact and material availability**

Further information and requests for resources and reagents should be directed to and will be fulfilled where possible by the lead contact, Amy Buck

## Data and code availability

The small RNA sequencing data are deposited to the National Center for Biotechnology Information (NCBI) Sequence Read Archive (SRA) under the BioProject ID PRJNA1200757. R scripts for analysis of the small RNA libraries are available at https://github.com/imu93/exwago_vaccine.

## Supplemental information

Document S1. Word file containing Figures S1-S4, Table S1, SI references, and legends for Datasets S1 and S2.

Dataset S1. Excel file containing data too large to fit in a PDF

Dataset S2. Excel file containing data too large to fit in a PDF, related to Figure 4A

