## Supplementary Information for "A non-vesicular Argonaute protein is transmitted from nematode to mouse and is important for parasite survival"

### denotes current affiliation

**This PDF file includes:**

Figures S1 to S4
Table S1
Legends for Datasets S1 and S2
SI References

**Other supporting materials for this manuscript include the following:**

Datasets S1 and S2

**Supplementary Figures**

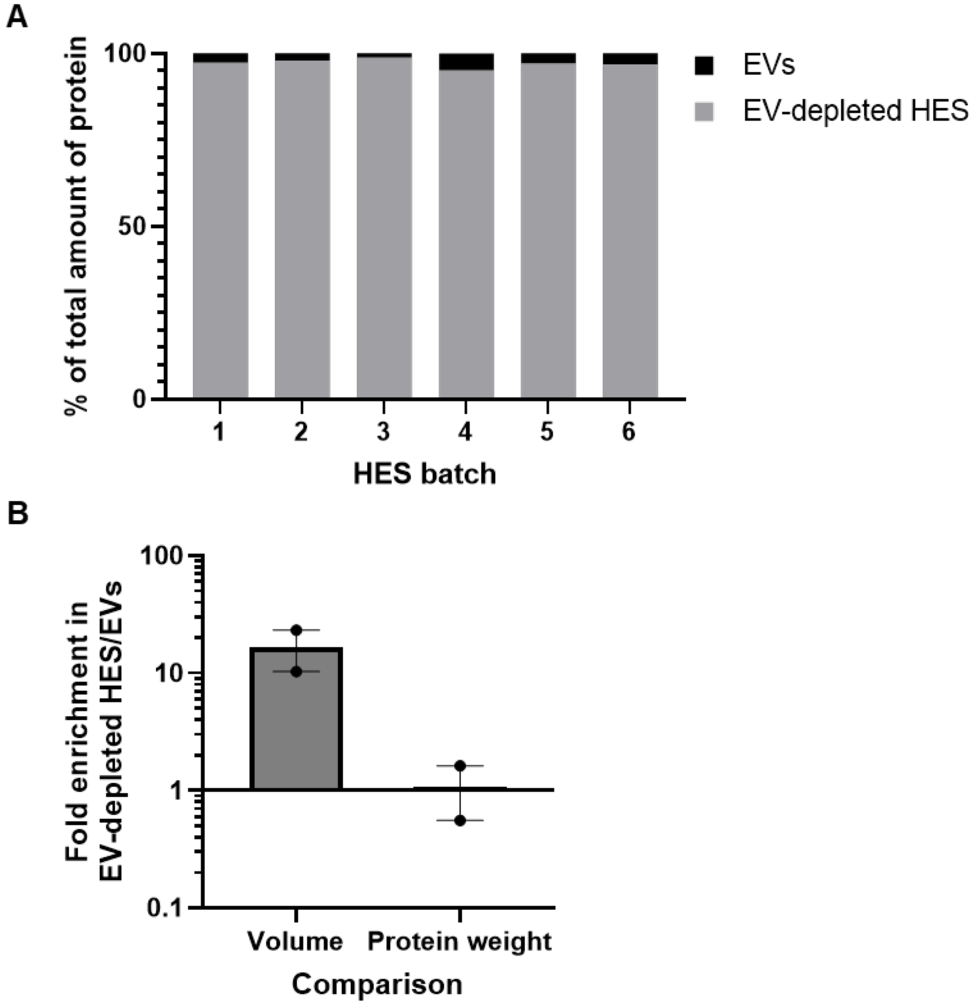


**Figure S1. Total protein yield of EV and EV-depleted HES and higher amounts of exWAGO in EV-depleted HES**

A) The amount of total protein quantified in EV-depleted HES and EVs following separation of HES by ultracentrifugation, shown as percentage of total protein of both fractions combined. Data are from 6 independent batches of HES collected from day 1-8 post-harvest of worms from mice. B) Fold-enrichment of Hb-exWAGO in EV-depleted HES compared to EVs following quantification of western blot band intensities. EV-depleted HES and EVs from the same starting material were analysed based on loading from equivalent starting sample volumes (left) or based on loading equivalent total protein (2.0 µg). Data represent mean ± S.E.M (n = 2 biological replicates).

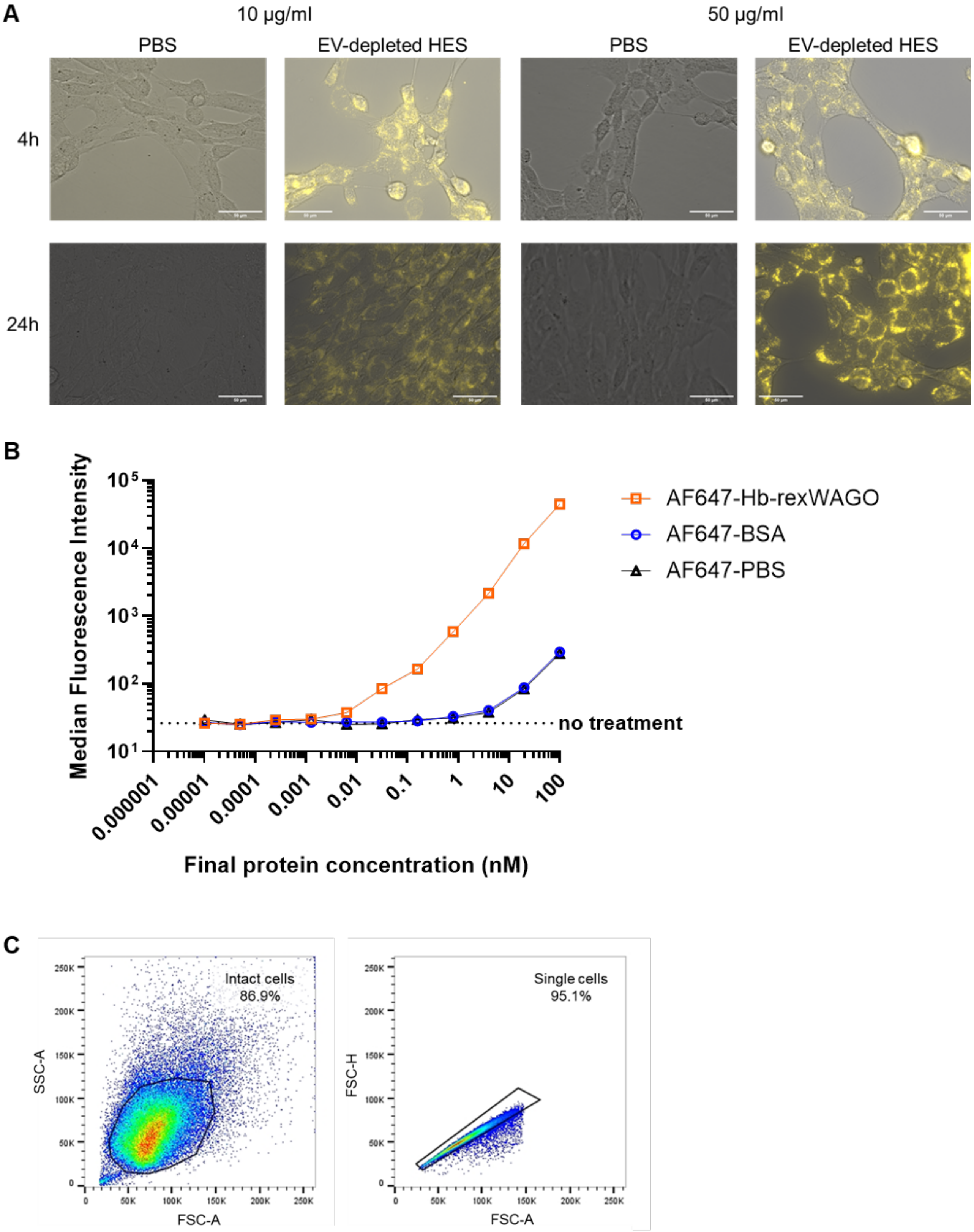
A
10 µg/ml
50 µg/ml
PBS
EV-depleted HES
PBS
EV-depleted HES
4h
24h
B
Median Fluorescence Intensity
$10^5$
$10^4$
$10^3$
$10^2$
$10^1$
no treatment
0.000001
0.00001
0.0001
0.001
0.01
0.1
1
10
100
Final protein concentration (nM)
AF647-Hb-rexWAGO
AF647-BSA
AF647-PBS
C
Intact cells
86.9%
SSC-A
FSC-A
Single cells
95.1%
FSC-H
FSC-A

**Figure S2. Detection of fluorescently-labelled proteins internalised by MODE-K cells**

A) Bright field and fluorescence composite microscopy images from MODE-K cells incubated with 10 or 50 µg/ml Cy-5-labelled EV-depleted HES or Cy-5-labelled PBS for 4 or 24 hours. Scale bar = 50 µm. Data are representative of 2 independent experiments. B) Dose response of AF647-labelled recombinant Hb-exWAGO (Hb-rexWAGO), BSA or PBS concentrations incubated with MODE-K cells for 4 hours measured by Flow Cytometry. C) Gating strategy applied for selecting intact cells (left panel) followed by selection of single cells (right panel) applied for Flow cytometry analysis.

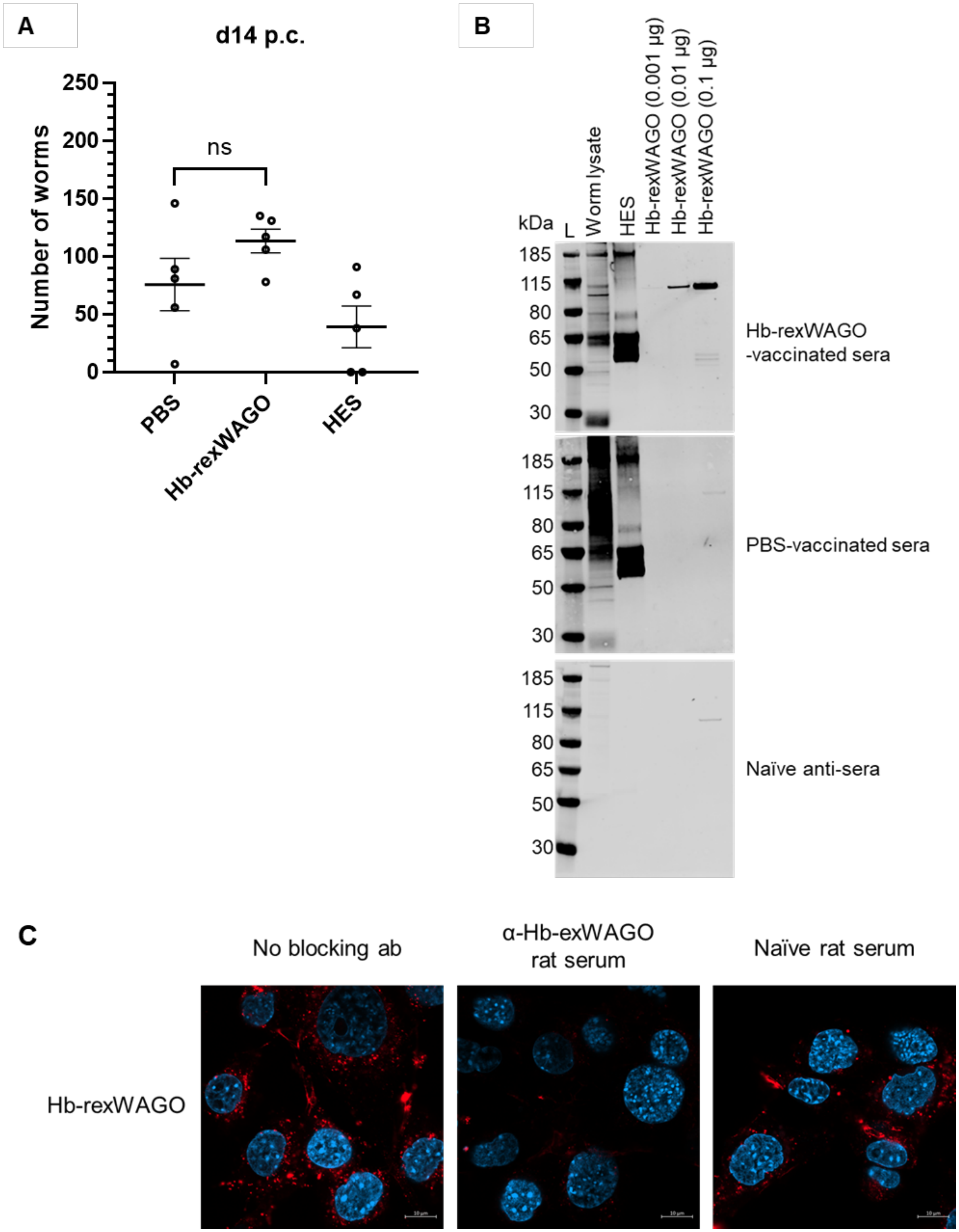


**Figure S3. Parasite and serological responses to vaccine**

A) The number of adult worms 14 days post-challenge (p.c.) recovered in the small intestine following vaccination of mice with recombinant Hb-exWAGO (Hb-rexWAGO),

HES, or PBS and challenge with 200 L3 stage larvae (n = 5 mice per vaccination group, from 1 experiment). Data represent the mean ± S.E.M. Data were analysed using an unpaired Kruskal-Wallis test (ns = $p > 0.05$). B) Western blot analysis of IgG responses using pooled sera from Hb-exWAGO- or PBS-vaccinated mice 28 days post-challenge (pool is from n = 5 mice in one experiment), or unvaccinated and uninfected mouse (referred to as naïve anti-sera, n = 1 mouse). Pooled sera were used as the primary antibody (1:1,000 in 5% BSA/TBST) and blots were probed with goat anti-mouse IgG AF680 antibody (1:10,000 in 5% BSA/TBST). Adult worm lysate = 1 µg; HES = 1 µg; recombinant Hb-exWAGO = 0.001-0.1 µg, L = ladder. C) Confocal super-resolution microscopy images of MODE-K cells incubated with 0.1 µM AF647-labelled Hb-rexWAGO for 4 hours in the presence of rat anti-exWAGO or naive serum (1:167 dilution). Data are representative of 3 independent experiments. Scale bar = 10 µm.

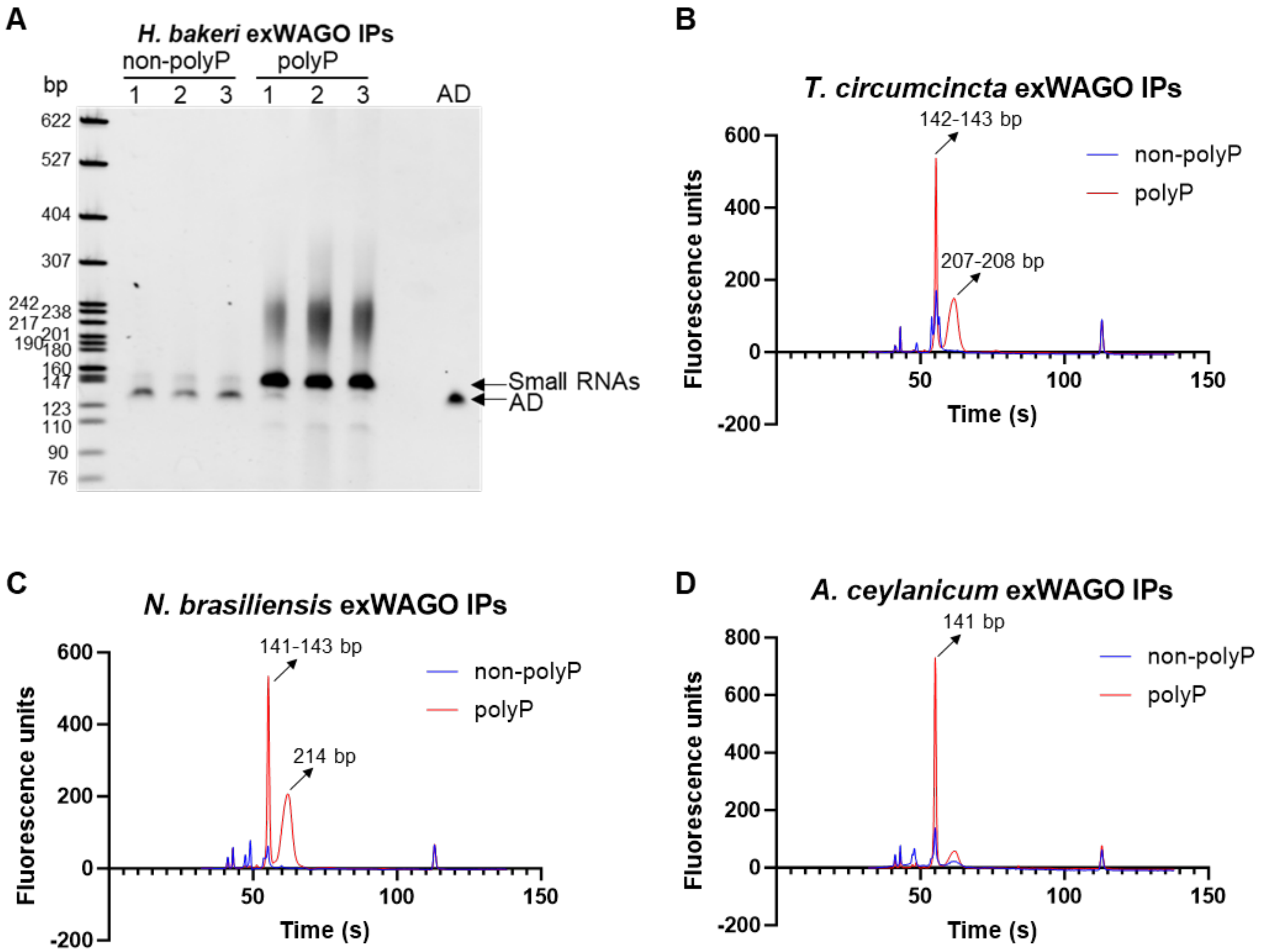


**Figure S4. Analysis of small RNA library profiles treated with or without 5' RNA polyphosphatase indicates exWAGO guides are 5' triphosphorylated**

A) Size profile of the small RNA libraries generated following Hb-exWAGO immunoprecipitation from *H. bakeri* adult worms prior to size selection based on TBE 10% gel. 1-3 denote biological replicates. AD = adapter dimers; non-polyP = non-polyphosphatase treated; polyP = polyphosphatase treated. B-D) Bioanalyser High Sensitivity DNA analysis of the small RNA libraries following exWAGO immunoprecipitation from adult B) *T. circumcincta*, C) *N. brasiliensis* and D) *A. ceylanicum* worms. The products enriched during polyphosphatase treatment are indicated. Non-polyP = non-polyphosphatase treated; polyP = polyphosphatase treated.

**Table S1**

Peptide sequences identified as the exWAGO orthologues in the excretory/secretory (ES) material of Clade V nematodes by mass spectrometry identified by Chow et al, (2019), Rooney et al, (2022), and Uzoechi et al, (2023). *Based on re-annotation of *T. circumcincta* genome (Dataset S1).

| Nematode species | Accession number | Length (aa) | Peptide sequence | Material | Reference |
|---|---|---|---|---|---|
| *Heligmosomoides bakeri* | HPOL_0000298601-mRNA-1 | 912 | TGMGQLSVGAVALPEKR<br>SAAVAVYK<br>AAVLFSAQR<br>QFMLPASVVSSAGPDATGIR<br>ISQMSIFFDQR<br>NAMQPFNQK<br>VTLQQQTPDQVASMIK<br>ASATLPQTR<br>IMKDALDITPR<br>AATTIAPR<br>LVNDGDLK | Adult EVs | (Chow *et al*, 2019) |
| *Nippostrongylus brasiliensis* | NBR_exWAGO | 913 | QDFVCNLTALK<br>DIFPQDSALFYDR<br>ILPTPTILYGER | Adult total ES | (Chow *et al*, 2019) |
| *Teladorsagia circumcincta* | TELCIR_10719 | 414 (950*) | TFPIGNLAAAPNALK<br>LFVGFEISNPPALSK | Adult EV-depleted ES | (Rooney *et al*, 2022) |
| *Ancylostoma ceylanicum* | ACEY_11634-1 | 912 | FQTTDGTQCTVEQYFK<br>NQNQTLDNVVAK<br>TFPIGNLAQPANQLK<br>VTDGFQVTSNDLQK | Female total ES | (Uzoechi *et al*, 2023) |

**Dataset S1 (separate file)**

The percentage amino acid identity of the exWAGO orthologues found in Clade V nematodes relative to Hb-exWAGO. Previously curated exWAGO protein annotations (Chow *et al*, 2019) were used to identify genomic regions and predicted protein products in new genome assemblies using miniport (Li, 2023).

**Dataset S2 (separate file)**

Alignment statistics and exWAGO expression quantification. RNA-seq libraries were aligned to their respective species assembly using STAR (Dobin *et al*, 2013). The "Aligned reads" column includes unique and multimapping reads. TPM (transcripts per million) values were calculated using overall gene counts, normalised by effective gene length.
